# Analytical results for directional and quadratic selection gradients for log-linear models of fitness functions

**DOI:** 10.1101/040618

**Authors:** Michael B. Morrissey, I. B. J. Goudie

**Affiliations:** School of Biology, University of St Andrews; School of Mathematics and Statistics, University of St Andrews

**Keywords:** natural selection, selection gradients, fitness, generalised linear model, capture-mark-recapture, survival analysis

## Abstract

1. Established methods for inference about selection gradients involve least-squares regression of fitness on phenotype. While these methods are simple and may generally be quite robust, they do not account well for distributions of fitness.
2. Some progress has previously been made in relating inferences about trait-fitness relationships from generalised linear models to selection gradients in the formal quantitative genetic sense. These approaches involve numerical calculation of average derivatives of relative fitness with respect to phenotype.
3. We present analytical results expressing selection gradients as functions of the coefficients of generalised linear models for fitness in terms of traits. The analytical results allow calculation of univariate and multivariate directional, quadratic, and correlational selection gradients from log-linear and log-quadratic models.
4. The results should be quite generally applicable in selection analysis. They apply to any generalised linear model with a log link function. Furthermore, we show how they apply to some situations including inference of selection from (molecular) paternity data, capture-mark-recapture analysis, and survival analysis. Finally, the results may bridge some gaps between typical approaches in empirical and theoretical studies of natural selection.

## 1 Introduction

The characterisation of natural selection, especially in the wild, has long been a major research theme in evolutionary ecology and evolutionary quantitative genetics (Endler, 1986; Kingsolver et al., 2001; Lande & Arnold, 1983; Manly, 1985; Weldon, 1901). In recent decades, regression-based approaches have been used to obtain direct selection gradients (especially following Lande & Arnold 1983), which represent the direct effects of traits on fitness. These, and related, measures of selection have an explicit justification in quantitative genetic theory (Lande, 1979; Lande & Arnold, 1983), which provides the basis for comparison among traits, taxa, etc., and ultimately allows meta-analysis (e.g., Kingsolver et al. 2001). Selection gradients can characterise both directional selection and aspects of non-linear selection, and so are a very powerful concept in evolutionary quantitative genetics.

Formally, the selection gradient is the vector of partial derivatives of relative fitness with respect to phenotype, averaged over the distribution of phenotype observed in a population. Given an arbitrary function ***W*(z)** for expected fitness of a (multivariate) phenotype **z**, a general expression for the directional selection gradient ***β*** is

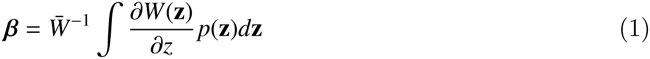

where *p*(**z**) is the probability density function of phenotype, and 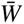 is mean fitness. Mean fitness can itself be obtained by ∫*W*(**z**)*p*(**z**)*d***z**. A quadratic selection gradient can also be defined as the average curvature (similarly standardised), rather than the average slope, of the relative fitness function,

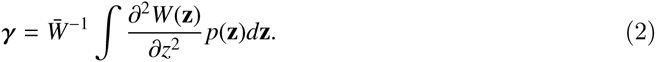

The directional selection gradient has a direct relationship to evolutionary change, assuming that breeding values (the additive genetic component of individual phenotype, Falconer 1960) are multivariate normally-distributed, following the Lande (1979) equation

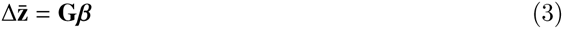

where 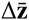 is per-generation evolutionary change, and **G** is the additive genetic covariance matrix, i.e., the (co)variances among individuals of breeding values. The quadratic selection gradient matrix has direct relationships to the change in the distribution of breeding values due to selection, but not with such simple relationships between generations as for the directional selection gradient and the change in the mean (Lande & Arnold, 1983).

The primary method for obtaining selection gradient estimates has been a simple and robust approach justified in Lande & Arnold (1983). The method involves least-squares multiple regression of relative fitness, i.e., absolute fitness divided by the mean observed in any comparable group of individuals over a specific period of the life cycle, potentially the entire life cycle, on measures of phenotype. Fitness, or any component of fitness, will typically have highly non-normal residuals in such a regression. Nonetheless, the simple least-squares methods are unbiased (see Geyer & Shaw 2010). However, methods that account for distributions of residuals that arise in regressions involving fitness as a response variable may provide better precision and more reliable statements about uncertainty (i.e., standard errors, p-values, etc.).

Some progress has been made at developing generalised regression model methods for inference of selection gradients. Janzen & Stern (1998) proposed a method for binomial responses (e.g., per-interval survival, mated vs. not mated). The Janzen & Stern (1998) method provides estimates of ***β***, and requires fitting a logistic model with linear terms only, calculating the average derivatives at each phenotypic value observed in a sample, and then standardising to the relative fitness scale. Morrissey & Sakrejda (2013) expanded Janzen & Stern’s (1998) basic approach to arbitrary fitness functions (i.e., not necessarily linear) and arbitrary response variable distributions, retaining the basic idea of numerically averaging the slope (and curvature) of the fitness function over the distribution of observed phenotype. Shaw & Geyer (2010) developed a framework for characterising the distributions of fitness (and fitness residuals) that arise in complex life cycles, and also showed how the method could be applied to estimate selection gradients by averaging the slope or curvature of the fitness function over the observed values of phenotype in a sample.

Of the many forms regression analyses of trait-fitness relationships might take, log-linear or log-quadratic models of the relationship between traits and expected absolute fitness may be particularly useful. In generalised linear models, the log link function is often useful and pragmatic. Fitness is a necessarily non-negative quantity, and expected fitness will typically best be modelled as a strictly positive quantity. This will indeed be the case if expected fitness is an exponential function of the sum of the predictors of the regression model, or, equivalently, a log link is used. Also, a log link function is compatible with generalised linear models with various distributions that could be useful for modelling fitness or fitness components. For example, it can be used with the Poisson distribution (counts, e.g., number of mates or offspring), the negative binomial distribution (for counts that are overdispersed relative to the Poisson distribution, potentially including lifetime production of offspring), and the exponential and geometric distributions (e.g., for continuous and discrete measures of longevity). The purpose of this short paper is to investigate the relationships between log-linear and log-quadratic models of fitness functions, and selection gradients.

## 2 Log-linear and log-quadratic fitness functions, and selection gradients

Selection gradients turn out to have very simple relationships to the coefficients of log-linear regression models predicting expected fitness from (potentially multivariate) phenotype. Suppose that there are k traits in the analysis and that the absolute fitness function, *W*(**z**) takes the form

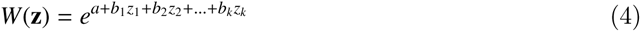

where *a* is a log-scale intercept, and the *b*_*i*_ are log-scale regression coefficients relating the traits (*z*_*i*_) to expected fitness. The equation for the directional selection gradient (equation 1) can then be simplified. Focusing on the selection gradient for a specific trait, i, in a log-linear model of *W*(**z**),

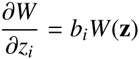

and hence

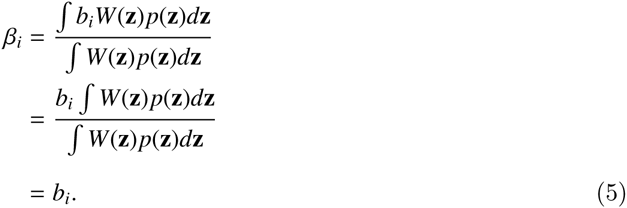

This result could be quite useful. In any log-linear model regressing expected absolute fitness, or a component of fitness, on trait values, the linear predictor-scale regression coefficients are the directional selection gradients.

The situation is a little bit more complicated if a log-quadratic model is fitted. If *W*(**z**) takes the form

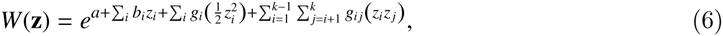

i.e., of a log-scale regression model with linear and quadratic terms, plus first-order interactions, then the b_i_ coefficients are not necessarily the directional selection gradients, nor are the *g*_*i*_ and *g*_*ij*_ coefficients the quadratic and correlational selection gradients, as they would be in a least squares analysis following Lande & Arnold (1983). However, we can use the log-scale quadratic fitness function with the general definitions of selection gradients (equations 1 and 2) to obtain analytical solutions for ***β*** and **γ**.

The factor of 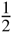 associated with the quadratic terms in equation 6 is a potential source of confusion, analogous to that surrounding a similar factor in Lande & Arnold’s (1983) paper (see Stinchcombe *et al.* 2008). In order to obtain the correct values of the *gi* coefficients, the covariate values for quadratic terms should be (1) mean-centred, then (2) squared, and then (3) halved. An alternative analysis is possible, where the squared covariate values are not halved, but the estimated coefficient estimates are doubled (analogous to procedures discussed by Stinchcombe *et al.* 2008). However, this alternative analysis leads to an additional, and potentially confusing, step in the calculation of standard errors (detailed in the appendix).

Define a vector **b** = (*b*_1_,’, *b*_k_)′ containing the coefficients of the linear terms in the exponent of the model in equation 6, and a matrix **g** = (*g_ij_*) containing the coefficients of the corresponding quadratic form. We can then write the fitness function more conveniently in matrix form

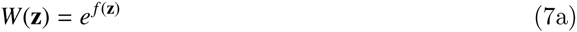

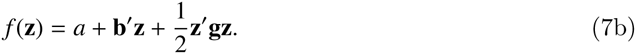

Let **d** be a vector of the expectations of the first order partial derivatives of *W*(**z**) and let **H** be the matrix of expectations of the second order partial derivatives of *W*(**z**). Thus the elements of **d** are 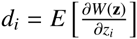 and the elements of **H** are 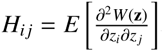. We can now rewrite the expressions for directional and quadratic selection gradients as

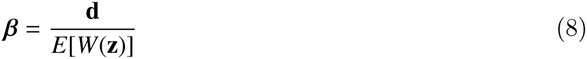

and

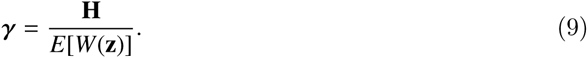

Differentiating equation 7 gives

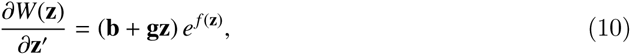

and

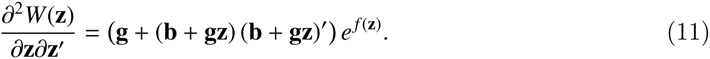

Assume that the phenotype **z** is multivariate normal, with mean ***μ*** and covariance matrix **Σ**, and denote its probability density by *p*_*μ*,Σ_(**z**). Provided *e*^*f*(**z**)^ has a finite expectation, the function

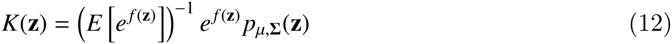

is a probability density function. Define the matrix **Ω**^−1^ = **Σ**^−1^-**g** and the vector *v* = *μ*+**Ω**(**b+gμ**). We show in the Appendix that **Ω** is symmetric. Provided it is also positive definite, it is a valid covariance matrix, and, by equation A7,

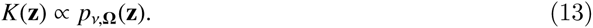

As *K* is a probability density function this implies,

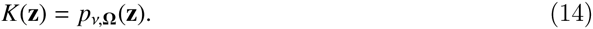

Define **Q**^−1^ = **Ω**^−1^Σ = **I**_k_-g**Σ**. Combining equations 8, 10 and 14 yields ***β*** = *E*[**b+gz**], where the expectation is taken with respect to *K*. This is an expectation of a linear function of **z**, and so

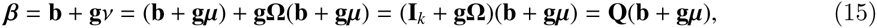

by use of equation A4.

Combining equations 9, 11 and 14 yields ***γ*** = *E*[**g + (b + gz) (b + gz)**’], where the expectation is taken with respect to *K*. Hence

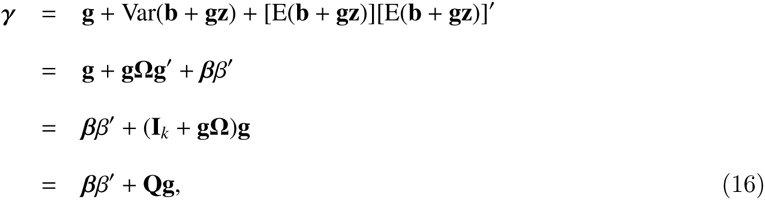

where we have noted that **g** is symmetric and used equation A4.

In univariate analyses, the matrix machinery necessary for implementing the general formulae in equations 15 and 16 can be avoided. If the fitness function is 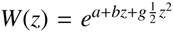 (note, again, that the quadratic coefficient is that for centred, then squared, and then halved values of z^1^), and *z* has a mean of *μ* and a variance of σ^2^ and then 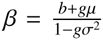 and 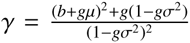. These expressions will hold for any univariate analysis, and can be applied to get mean-standardised, variance-standardised, and unstandardised selection gradients, when appropriate values of and σ^2^ are used, and applied to log-quadratic models of *W*(***z***) where the phenotypic records have been correspondingly standardised. For the common case where the trait is mean-centred and (unit) variance standardised, the expressions simplify further to 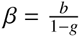 and 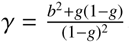

The equivalence of the regression coefficients of a log-linear fitness model with directional selection gradients (equation 5) of course requires that the regression model provides a reasonable description of the relationship between a trait and expected fitness and makes reasonable assumptions about fitness residuals. Otherwise, the relationship is relatively unburdened by assumptions. For example, it does not require any specific distribution of phenotype. The use of selection gradients obtained from log-linear regressions to predict evolution using the Lande equation (equation 3) does assume that breeding values are multivariate normal (see Morrissey 2014 for a discussion of selection gradients and associated assumptions about multivariate normality of phenotype and breeding values). The expressions for ***β*** and ***γ*** given a log-quadratic fitness model (equations 15 and 16) do assume multivariate normality of phenotype. Equations 15 and 16 further require that **Ω** is positive definite. In univariate analyses, this condition reduces to 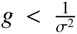, implying that the fitness function should not curve upwards too sharply within the range of observed phenotype.

A very convenient feature of the expressions for ***β*** and ***γ*** in equations 5, 15 and 16 is that the model (log) intercept does not influence the selection gradients. This means that the range of modelling techniques that yield selection gradients can be even further expanded. For example, adding fixed and random effects to Lande & Arnold’s (1983) least squares analysis will generally result in estimated regression coefficients that are not interpretable as selection gradients. For example, it might be desirable to estimate a single selection gradient across two sexes, if data are limited and sex-differences in selection are not anticipated. In such an analysis, it might seem sensible to include an effect of sex, to account for differences in mean fitness between the sexes. However, such an analysis would not yield correct selection gradients, because the theory underlying the least squares-based regression analysis of selection requires that mean relative fitness is one, and this would not be the case when different strata within an analysis have different intercepts. On the other hand, adding such an effect to a log-scale model of absolute fitness, and then deriving selection gradients using equations 5, 15 and 16 will yield correct selection gradients. Other effects, such as random effects to account for individual heterogeneity in expected fitness, beyond that explained by the traits (or correlated, unmeasured traits), will be usable as well, while still retaining the ability to obtain correct selection gradients.

## 3 Statistical uncertainty

The expressions for selection gradients, given the parameters of a log-quadratic fitness function (equations 15 and 16) give the selection gradients conditional on the estimated values of **b** and **g**. However, **b** and **g** will not typically be known quantities in empirical studies of natural selection, but rather will be estimates with error. Because equations 15 and 16 are non-linear functions of one or more regression coefficients, unconditional estimators of ***β*** and ***γ*** would have to be obtained by integrating the expressions for ***β*** and ***γ*** over the sampling distributions of the estimated values of **b** and **g**. Such details are not normally considered in calculations of derived parameters (e.g., heritabilities) in evolutionary studies. Such integration could be achieved using approximations, bootstrapping, or MCMC methods. Alternatively, application of equations 15 and 16 directly to estimated values of b and g may be sufficient in practice. Similarly, while standard errors of the parameters b and g are not directly interpretable as standard errors of corresponding values of ***β*** and ****β****, approximations, bootstrapping, and MCMC methods may all potentially be useful in practice. In particular, approximation of standard errors by a first-order Taylor approximation (the “delta method”; Lynch & Walsh 1998) may generally be pragmatic. Formulae for approximate standard errors by this method are given in the appendix. For univariate analysis, with phenotype standardised to *μ* = 0 and *σ*^2^ = 1, the approximate standard errors of *β* and *γ* are given by

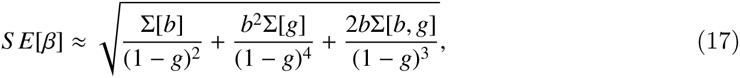

and

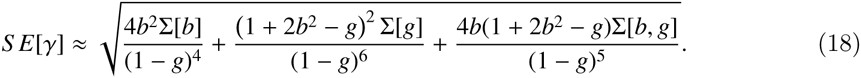

Where Σ[b] and Σ[g] represent the sampling variances of the estimated *b* and *g* terms. These are the squares of their standard errors. Σ[*b, g*] is the sampling covariance of the *b* and *g* terms. This is not always reported, but can usually be obtained. For example, in R, it can be extracted from a fitted glm object using the function vcov().

We performed a small simulation study to assess the extent of any bias in the estimators ***β*** and ***γ*** and the adequacy of their standard errors. We simulated univariate directional selection, with values of *b* between −0.5 and 0.5, and with *g* = −0.5,0 and 0.2. Because *β* and *γ* are nonlinear functions of *g*, it is not possible to simultaneously investigate ranges of parameter values with regular intervals of values of both *g* and selection gradients. These values of *g* represent a compromise between investigating a regular range of *g* and *γ*. We used a (log) intercept of the fitness function of *a* = 0. We simulated a sample size of 200 individuals. This sample size reflects a very modest-sized study with respect to precision in inference of non-linear selection, and is therefore a useful scenario in which to judge performance of different methods for calculating standard errors. Fitness was simulated as a Poisson variable with expectations defined by the ranges of values of *b* and *g*, and with phenotypes sampled from a standard normal distribution.

Firstly we analysed each simulated dataset using the OLS regression described by Lande & Arnold (1983), i.e., 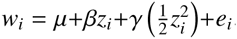, using the R function lm(). For the OLS regressions, we calculated standard errors assuming normality using the standard method implemented in the R function summary. lm (), and by case-bootstrapping, by generating 1000 bootstrapped datasets by sampling with replacement, running the OLS regression analysis, and calculating the standard deviation of the bootstrapped selection gradient estimates. Secondly we fitted a Poisson glm with a linear and quadratic terms, using the R function glm(). We then calculated conditional selection gradient estimates using equations 15 and 16. We obtained standard errors by using a first-order Taylor series approximation (the “delta method”; Lynch & Walsh 1998, appendix A1). For each method of obtaining estimates and standard errors, we calculated the standard deviation of replicate simulated estimates. We could thus evaluate the performance of different methods of obtaining standard errors by their ability to reflect this sampling standard deviation. We also calculated mean absolute errors for both estimators of *β* and *γ* for all scenarios. Every simulation scenario and associated analysis of selection gradients was repeated 1000 times.

Selection gradient estimates obtained by all three methods were essentially unbiased (figure 1a,d,g,j,m,p), except for small biases that occurred when the fitness function was very curved. Thus, glm-derived values of selection gradients, conditional on estimated values of *b* and *g* performed very well as estimators of *β* and *γ* in our simulations. Similarly, first-order approximations of standard errors of the glm-derived estimates of *β* and *γ* closely reflected the simulated standard deviations of the estimators (figure 1). All methods for obtaining standard errors performed well for estimates of *β* in the pure log-linear selection simulations (figure 1h,k). OLS standard errors performed reasonably well under most simulation scenarios, except when g was positive (figure 1n,q); across all scenarios bootstrap standard errors of the OLS estimators outperformed standard OLS standard errors. Mean absolute error of the glm estimators was always smaller than that of the OLS estimators of *β* and *γ*. This is unsurprising, as the simulation scheme corresponded closely to the glm model. These results demonstrate the usefulness of the conditional values of *β* and γ as estimators, and show that gains in precision and accuracy can be obtained when glm models of fitness functions fit the data well. It remains plausible that the OLS estimators motivated by Lande & Arnold’s (1983) work could outperform glm-based analyses in some scenarios.

**Figure 1:**
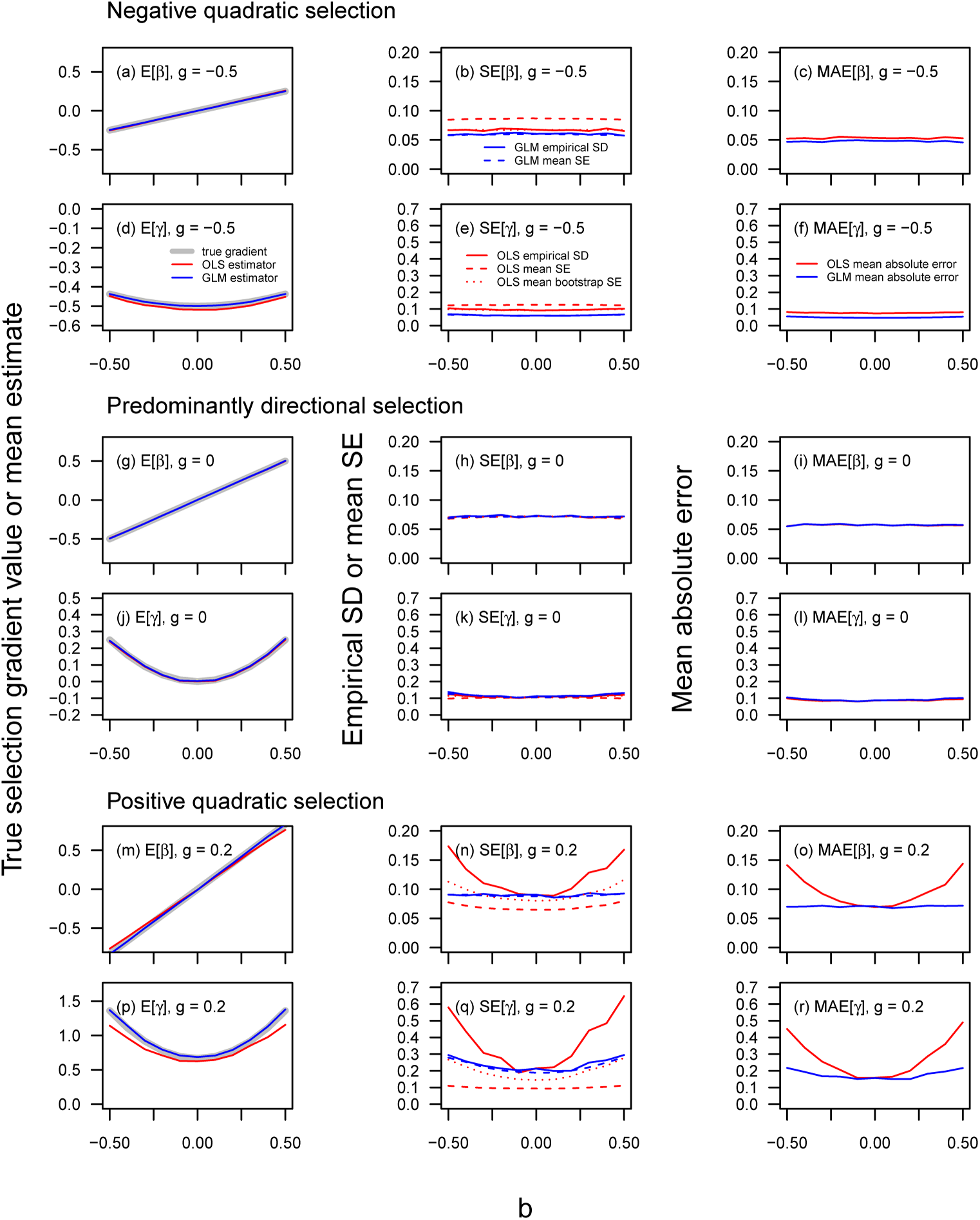
Simulation results for the performance of Lande & Arnold’s (1983) least squares-based (OLS) estimators (red lines), and log-quadratic (GLM) estimators (blue lines), of directional and quadratic selection gradients. The first column shows bias in estimates of *β* and *γ*, where departure from the grey line (the simulated truth) indicates bias. The middle column shows the performance of OLS standard errors (red dashed lines), bootstrap standard errors (red dotted lines), and first-order approximations (blue dashed lines) of the standard errors of the GLM estimators. Ideally, all values of estimated mean standard errors would fall on the simulated standard deviation of their associated estimators, shown as solid lines. The right column shows the mean absolute errors of the OLS and GLM estimators.

## 4 Other analyses that correspond to log-linear fitness functions

In addition to generalised linear models with log link functions, there may be other cases where models of trait-fitness relationships may correspond to log-linear or log-quadratic fitness functions. In paternity inference, some methods have been proposed wherein the probability that candidate father i is the father of a given offspring is modelled according to

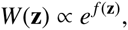

and where realised paternities of a given offspring array are then modelled according to a multinomial distribution, potentially integrating over uncertainty in paternity assignments based on molecular data (Hadfield et al., 2006; Smouse et al., 1999). When f(z) is a linear function, Smouse, Meagher & Korbak (1999; T. Meagher, personal communication) interpreted the analysis as analogous to Lande and Arnold’s 1983, but not necessarily identical. For a linear f (z), this analysis does in fact yield estimates of ***β***, and for a quadratic function, directional and quadratic selection gradients can be obtained using equations 15 and 16. This can be seen by noting that expected fitness, given phenotype, of candidate fathers for any given offspring array will be, in the log-linear case,

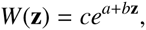

where *c* is a constant. In application of the expressions yielding equation 5, c appears in both the numerator and the denominator, yielding ***β*** = **b**.

Another case where our formulae may be applicable pertains to inferences of survival rate. Often, data about trait-dependent survival rates may be assessed over discrete intervals. While the experimental unit of time may be an interval (e.g., a day or a year), the biologically-relevant aspect of variation in survival may be longevity, i.e., for how many intervals an individual survives. One such situation arises when per-interval survival rate is assessed via a logistic regression analysis, and trait-dependent survival rates are (or may be assumed to be) constant across intervals. A common case of logistic regression analysis that satisfies this first condition is often implemented in capture-mark-recapture procedures. Suppose that per-interval survival rate, given phenotype, may be assumed to be constant, and that fitness is defined to be the expected survival time. Then fitness will be given by the mean of a geometric distribution where death in a particular interval of an individual with phenotype z occurs with probability ρ(**z**),

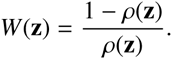

If trait-dependent per-interval survival probability is denoted *φ*(**z**) (*φ* being the standard symbol for survival rate in capture-mark-recapture analyses; Lebreton et al. 1992), then the fitness function in terms of expected number of intervals lived is 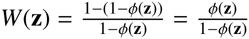. If per-interval survival rate has been modelled as a logistic regression, i.e.,

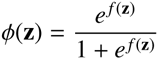

where φ(**z**) denotes the per-interval fitness function, and *f*(**z**) is the fitness function on the logistic scale, then the fitness function on the discrete longevity scale is

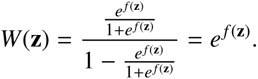

Therefore, if *f*(**z**) is a linear function, then its terms are the directional selection gradients on the discrete-longevity scale. If *f*(**z**) is a quadratic function, then the corresponding directional and quadratic selection gradients, again if the relevant aspect of fitness is the number of intervals survived, can be obtained using equations 15 and 16. Waller and Svensson (2016; this issue) takes advantage of these relationships to compare inference of trait-dependent survival in capture-mark-recpature models to classical inference using Lande & Arnold’s (1983) least-squares regression analysis where fitness is assessed as the number of intervals that individuals survive.

It must be stressed that these results do not justify interpretation of logistic regression coefficients of survival probability as selection gradients in a general way. Such coefficients differ from selection gradients for three reasons: (1) they pertain to a linear predictor scale, and natural selection plays out on the data scale, (2) they directly model absolute fitness, not relative fitness, and (3) they pertain to per-interval survival, which may not necessarily be the aspect of survival that best reflects fitness in any given study. It is only when the number of intervals survived is of interest (and mean survival can be assumed to be constant across intervals) that these three different aspects of scale cancel out such that the parameters of a logistic regression are selection gradients.

Finally, another situation where an important analysis for understanding trait-fitness relationships that has an immediate – but not necessarily immediately apparent – relationship to selection gradients, arises in survival analysis. In a proportional hazards model (Cox, 1972), the instantaneous probability of mortality experienced by live individuals, the hazard λ(t), as a function of their phenotype could be modelled as

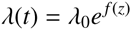

where λ_0_ is the baseline hazard, and the *e^f(z)^* part of the function describes individual deviations from this baseline hazard. If the baseline hazard is constant in time, then survival distributions conditional on phenotype are exponential, and have mean λ^−1^. So, if fitness is taken to be expected longevity (as a continuous variable now, not discrete number of intervals as in the relations given above between logistic models of per-interval survival and selection gradients) then

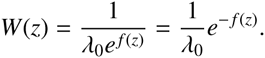

In expressions for selection gradients (equations 1 and 2), 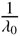 would be a constant in the integrals in both the numerators and denominators, and therefore cancels in calculations of selection gradients. Therefore, if proportional hazards are modelled with *f(z)* as a linear or quadratic function, then the expressions for selection gradients (equations 5, 15 and 16) hold, but the coefficients of the trait-dependent hazard function must be multiplied by −1.

## 5 Conclusion

We have provided analytical expressions for selection gradients, given the parameters of log-linear and log-quadratic functions describing expected fitness. These functions can be applied in conjunction with a range of generalised linear model approaches, specific situations in capture-mark-recapture analysis, and relate to fitness functions used in theoretical studies. The general relationship of selection gradients to the coefficients of log-linear and log-quadratic models, in particular, various generalised linear models, are probably the most generally useful feature of our results. In empirical applications, our preliminary simulation results indicate that, given an appropriate model of a log-scale fitness function, inference using log-linear and log-quadratic models may be very robust, and could provide more reliable statements about uncertainty (e.g., reasonable standard errors) than the main methods used to date. Furthermore, the relationships given here between log-quadratic fitness functions and selection gradients could lead to better integration between empirical and theoretical strategies for modelling selection. In theoretical studies, Gaussian fitness functions are often used. These are simply log-quadratic functions that are parameterised in terms of a location parameter (phenotype of maximum fitness), and a width parameter. A relationship between the parameters of a Gaussian fitness function and directional selection gradients (Lande 1979; the expression we give for *β* is an alternative formulation) is already widely used in the theoretical literature. For any given distribution of phenotype, these parameters correspond directly to linear and quadratic (log-scale) regression parameters, and so can be directly related to selection gradients in empirical studies.

## Acknowledgements

We thank Andy Gardner, Graeme Ruxton, and Kerry Johnson for discussions, comments, and advice. Peter Jupp provided particular insights that improved this paper. MBM is supported by a Royal Society (London) University Research Fellowship.

## Appendix

Denote a vector containing all unique elements of ***γ*** by 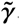 The following assumes that 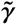 is composed by vertically stacking the columns of the diagonal and sub-diagonal elements of *γ.* For example, in an analysis with three traits, 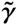 = γ_1,1_,γ_2,1_,γ_3,1_,γ_2,2_,γ_3,2_,γ_3,3_. Let **v**() denote the function mapping the distinct elements of a symmetric matrix **r** onto the column vector 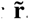.

The first-order approximation to the sampling covariance matrix of the elements of ***β*** and ***γ*** is then given by 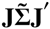, where 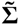 is the sampling covariance matrix of a vector containing the elements of **b** and 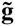, where the latter is a column vector containing the distinct elements of **g** arranged according to the same scheme that defines 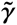. **J** is the Jacobian, or gradient matrix of first order partial derivatives, of ***β*** and 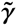 with respect to **b** and 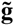, i.e.,

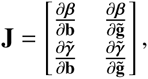

evaluated at the estimated values of **b** and **g**.

Note that some users may prefer to fit the model 6 with *g*_*ii*_ replaced by *2g*_*i*_, say. The formulae for ***β*** and ***γ*** are readily re-expressed in terms of these variables by making this substitution. If **Σ**_1_ denotes the covariance matrix obtained when fitting this revised model, the required covariance matrix 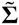 can be calculated using 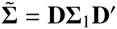, where **D** is a diagonal matrix with all the diagonal elements equal to one, apart from those corresponding to the variables *g*_ii_ which equal 2.

The four submatrices of **J** can be treated separately. Noting that ***β*** = **Q (b + g*μ*)** (equation 15),

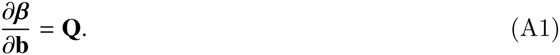

Let 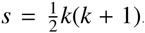, where *k* is the number of traits in the analysis, and let **e**_1_,…, **e**_s_ be the standard basis for an *s* dimensional space (i.e., **e**1 = [1,0,…,0]’, etc.). Define an indicator matrix **C**_m_ = **C**^*(i,j)*^ where **C**^*(i,j)*^ is a *k* by *k* matrix in which

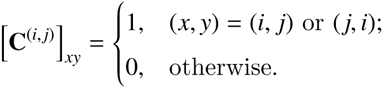

Using the standard expression for the derivative of the inverse of a matrix with respect to a scalar, we can obtain 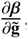, i.e., the upper-right sub-matrix of **J**.

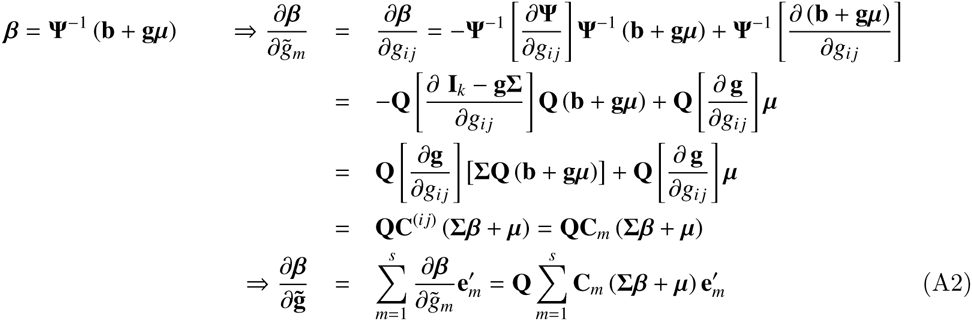

Let **Q**_[u]_ denote the *u*^th^ column of **Q**. Using the previous relation 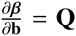, we can obtain 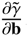, i.e., the lower-left sub-matrix of **J**.

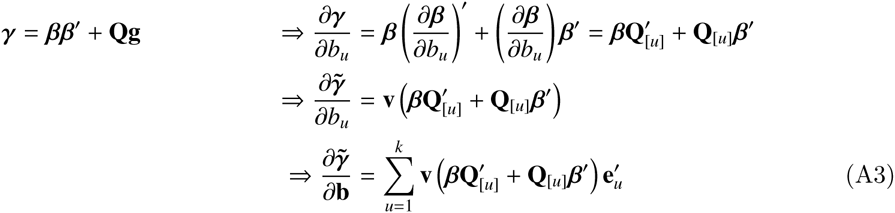

Let **M**^(m)^ = **QC**_m_ (**Σ***β*+*μ*)***β***′ Note that **Q**^−1^ = **Ω**^−1^Σ implies **Ω = EQ**. Moreover **Ω**^−1^ = **Σ**^−1^ implies firstly that

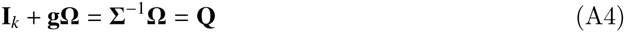

and secondly that **Ω** is symmetric, since **Σ** and **g** are both symmetric. It follows that

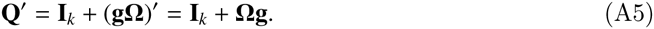

The lower-right sub-matrix of **J** can then be derived.

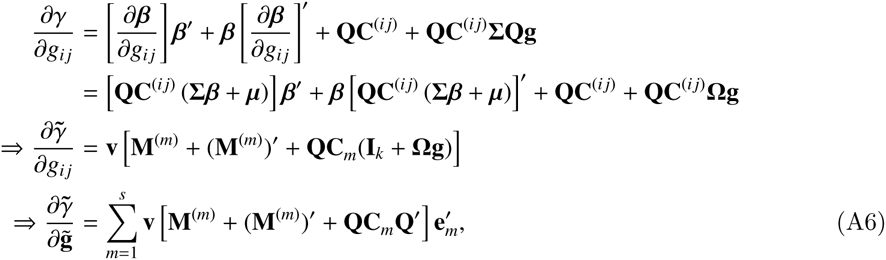

by use of equation A5.

Finally note that equations A4 and A5 are also relevant to the derivation of formula 13. By definition, 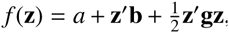, and we have log 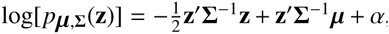 where *α* does not depend on **z**. Thus, if *α*′ = *α* + *a*, it follows that, as a function of **z**,

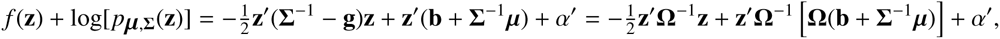

Now, by A4 and A5, we have **Ω(b+Σ^−1^*μ*)=Ωb + (Σ^−1^Ω)′*μ*) = Ωb + Q′*μ* = Ωb + (I_k_ + Ωg)*μ*** = v, implying that

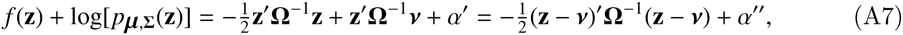

where *α″* is constant as a function of **z**. The exponent of *e*^*f*(**z**)^*p*_*μ*Σ(**z**)_ is thus identical, as a function of **z**, to that of *p*_vΩ(**z**)_. Hence formula 13 holds.

## Footnotes

1 This can be accomplished easily in R. Assume that *W* and *z* are variables in memory representing absolute fitness and phenotypic data, and that residuals of *W* are assumed to follow a Poisson distribution. The regression could be implemented by glm(W˜z+I(0.5*(z-mean(z))^2), family=poisson(link=“log”)).

